# Embryonic development of the Mediterranean starfish *Hacelia attenuata*

**DOI:** 10.64898/2025.12.04.692310

**Authors:** Silvia Caballero-Mancebo, Laurent Gilletta, Janet Chenevert, Stefania Castagnetti

**Affiliations:** Sorbonne Université, CNRS, Laboratoire de Biologie du Développemend de Villefranche-sur-mer (LBDV), 06230 Villefranche-sur-mer, France

## Abstract

Starfish play essential ecological roles as predators and ecosystem regulators, however detailed developmental descriptions exist for only a handful of species, none of which come from the Mediterranean. In this study we provide the first full account of the development of the Mediterranean starfish *Hacelia attenuata*, from oocyte maturation through embryogenesis and larval formation, showing that its development largely follows the canonical pattern known from other asteroids. Oocytes resume meiosis when exposed to 1-methyladenine, exhibiting conserved features such as formation of a nuclear actin shell for chromosome gathering prior to meiotic spindle assembly. Embryogenesis then proceeds through equal radial cleavages to form a ciliated blastula, followed by gastrulation via invagination and mesenchymal cell delamination. The larvae develop through typical bipinnaria and brachiolaria stages, displaying characteristic feeding structures and attachment organs. Importantly, developmental rate increases substantially at higher temperatures, consistent with the species distribution in warm Mediterranean waters. Together, these findings position *H. attenuata* as a promising new model for studying thermal adaptation, environmental resilience and conserved developmental mechanisms in starfish.

## Introduction

Starfish are marine animals found across all marine ecosystems, from tropical coral reefs to the deep seafloor. Most species are predators, feeding on various types of prey. By regulating the populations of organisms they feed on, starfish help prevent single species from dominating a habitat and shape the overall ecosystem structure thereby acting as keystone species (Burt et al., 2018; Paine, 1966). A classic exemple is *Asterias rubens*, a key predator of *Mytilus edulis* on North Atlantic tidal flats. It prevents mussel beds from becoming overly dense, thus promoting species diversity and allowing other invertebrates and algae to persist in the community (Dayton, 1971). Another well-known case is the crown-of-thorns starfish (*Acanthaster* spp.), a coral-eating species that primarily feeds on fast-growing branching and tabular corals, especially *Acropora*. In balanced populations, *Acanthaster* helps maintain coral diversity by limiting the dominance of rapidly growing corals, allowing slower-growing species to thrive. However, *Acanthaster* population outbreaks can lead to massive coral mortality and structural degradation of reefs, while their absence can result in coral monocultures and reduced reef resilience (Pratchett et al., 2009).

Starfish (Asteroidea) are ambulacrarians and, together with brittle stars (Ophiuroidea), form the subphylum Asterozoa within Echinodermata (Figure 1A). There are about 1900 known extant species of starfish worldwide which are classified into five major orders: Paxillosida, Spinulosida, Velatida, Valvatida and Forcipulata (Mah and Blake, 2012) and 37 families (Mah and Blake, 2012; Feuda and Smith, 2015; Byrne et al., 2017; Linchangco et al., 2017). Despite their diversity, detailed descriptions of starfish development exist only for a limited number of species. The most common laboratory-used species are *Patiria miniata*, which inhabits mostly the northeastern Pacific coastline, and *Patiria pectinifera*, commonly known as the blue bat star, native to the temperate northern Pacific region along the coasts of Japan, China, and Russia. Both species belong to the Valvatida order (Mah and Foltz, 2011). Besides these, development has been docucumented for a few additional species, such as the common North Atlantic sea star *Asteria rubens* (Carter et al., 2021) belonging to Forcipulata, and the crown of thorns *Acanthaster planci (*Lucas 1973). To our knowledge, however, no complete developoment has been yet reported for any Mediterranean starfish. Studies on egg maturation and fertilization have been carried out in few Mediterranean species, such as *Marthasterias glacialis* (Forcipulata, Fisher et al., 1998), *Echinaster sepositus* (Spinulosida, Picard et al., 1985) and *Astropecten aranciacus* (Paxillosida, Fisher et al., 1998), but their development remains undescribed.

**Figure 1:**
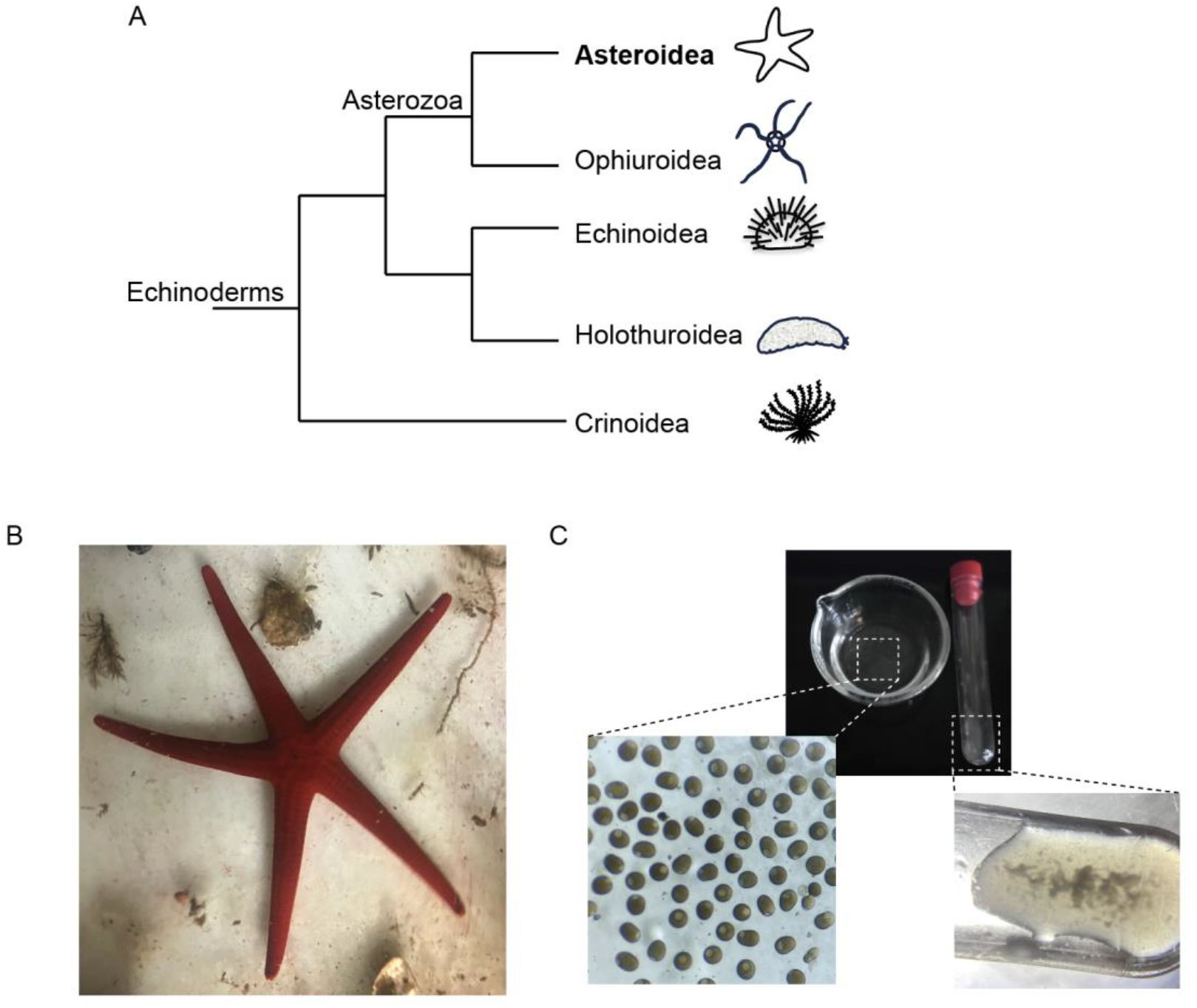
Gamete collection from the Mediterranean starfish *H. attenuata*. A) Phylogenetic tree of echinoderms with Asteroidea in bold. B) Image of adult specimen of *H. attenuata*, collected in the Bay of Villefranche. C) Images of gamates following collection from the arm of the animal.

Temperature, salinity, pH, and food availability strongly affect starfish larval survival, development and metamorphic success (Byrne, 2011). As climate change and ocean acidification intensify, these conditions are shifting, disrupting starfish population dynamics, undermining their ecological functions and potentially destabilizing entire marine ecosystems. Therefore, studying starfish development across oceans can reveal how environmental factors drive biological adaptations and shape biological traits. In this context, establishing a Mediterranean starfish model is especially valuable as it will help elucidate adaptations to environments with warmer, more stable temperatures, higher salinity, and relatively nutrient-poor waters compared with many open-ocean habitats.

Although the species was first described by John Edward Gray in 1840 (Gray, 1840), here we provide the first comprehensive description of the development of the smooth Mediterranea starfish *Hacelia attenuata* (Figure 1B) from egg maturation to larval development. By documenting the timing and morphology of key developmental stages under controlled laboratory conditions, and by comparing these features with those of the established model species *Patiria miniata*, we establish *H. attenuata* as a new model system for developmental biology.

## Matherial and methods

### Sampling and culturing of *H. attenuata*

Adult specimens of *H. attenuata* were collected in the Bay of Villefranche (France) at 30-40 meters depth throughout the year and maintained in acquaria at the Biological Resource Center in Villefranche (CRB) at 14°C. Adults were fed fresh mussels (*Mitylus galloprovincialis*) and could be maintained gravid for up to 10 months.

Fertilization and embryo rearing were performed in 0.22µm microfiltered sea water (MFSW) at 18 °C, 21 °C or 23 °C, in glass dishes. From opening of the mouth (early bipinnaria stage), larvae were fed daily with a mix of *Chaetoceros calcitrans* and *Tisochrysis luthea* (produced at CRB) at 8000-12000 cells/ml (1:1). Seawater was changed every day and larvae were moved to clean glass dishes once per week to avoid bacterial growth and to ensure proper development.

### Immunofluorescence and phalloidin staining

For immunofluorescence, eggs and embryos were fixed overnight in 90% methanol containing 50 mM EGTA at −20 °C. After fixation samples were washed 3 times in phosphate-buffered saline (PBS) containing 0.1% Tween-20 (PBST), then blocked in PBS containing 3% BSA for 1 h at room temperature, and finally incubated overnight at 4 °C in PBS containing 3% BSA and mouse anti-tubulin DM1A (Sigma-Aldrich) antibody diluted 1:500. Following 3 washes in PBST, eggs were incubated with anti-mouse fluorescently-labeled secondary antibody (1:500) at room temperature for 1–2 h. Following a wash in PBST, samples were further incubated for 10 min in PBST containing Hoechst (10 µg/mL), washed twice and then mounted in Citifluor AF1 (Science Services, München, Germany).

For phalloidin staining, eggs and embryos were fixed overnight in 3.7% paraformaldehyde at 4 °C. After fixation samples were washed three times in PBS and then permiabilized by incubation in PBS containing 0.1% Tween-20 (PBST) for 10 minutes. Samples were finally incubated overnight at 4 °C in 500 µl of PBST containing 2 unit of rhodamine phalloidin (Invitrogen; stock solution 1U/µl in dmso). Samples were washed once in PBS and then incubated in PBST containing Hoechst (10 µg/ml) for 30 minutes and then mounted directly in Citifluor AF1.

Imaging of fixed samples was performed either on a Leica SP8 confocal microscope or on a Leica Stellaris microscope equipped with a LASX software.

### Imaging and Time-Lapse Microscopy

For imaging live eggs, embryos and larvae, samples were mounted between slide and coverslip using Dow Corning vacuum grease as a spacer as described elsewhere [18] or were placed in dishes containing sea water. Images were acquired using a Zeiss Axioimager A2 upright microscope with differential interference contrast (DIC) optics and a AxioCam506 camara and Zen acquisition software.

For timelapse imaging, fertilized eggs were placed in glass bottom dishes (MatTek corporation, Ashland, MA, USA) containing MFSW and images were acquired every 1–2 min with 20× objective lenses on a Zeiss Axiovert 200 inverted microscope equipped with a Hamamatsu ORCA Fusion camera and a Metamorph acquisition software.

Alternatively, fertilized injected zygotes were mounted in multiwell sample holders and placed in the imaging chamber on a LS2 Live light sheet microscope from Viventis equipped with two Nikon CFI Plan Fluor 10x/0.2 NA objectives for illumination and two Nikon CFI75 Apochromat 25x/1.1 NA water immersion objectives. Detection was performed with two Hamamatsu ORCA Fusion cameras. Embryos were imaged *in toto* with an optical section of 2 μm every 10 min.

## Results

### Gametes collection meiotic progression and fertilization

*H. attenuata* adults were collected in the bay of Villefranche-sur-mer (France) at different times during the year. Animals collected between November and April contained little to no gamets, whereas animals collected from June to October, when sea water temperature rised above 16°C, had full gonads. Eggs and sperm could be easily collected by aspiration from the gonads, located in the arms of the animals, using a syringe with an 18G needle (Figure 1C), reducing the risk of infection compared to commonly used dissection. Upon collection, *H. attenuata* eggs were transparent and spherical, with a diameter of 154 +/-5 µm (Figure 2 A,B) and were arrested at prophase of meiosis I with a large nucleus, the germinal vesicle (GV), 69 +/-4 µm in diameter (Figure 2 A,B). The GV is off-centered, located near the surface of the egg and chromosomes are condensed and scattered in the entire space, often in the vicinity of the nuclear envelope (Figure 2C).

**Figure 2:**
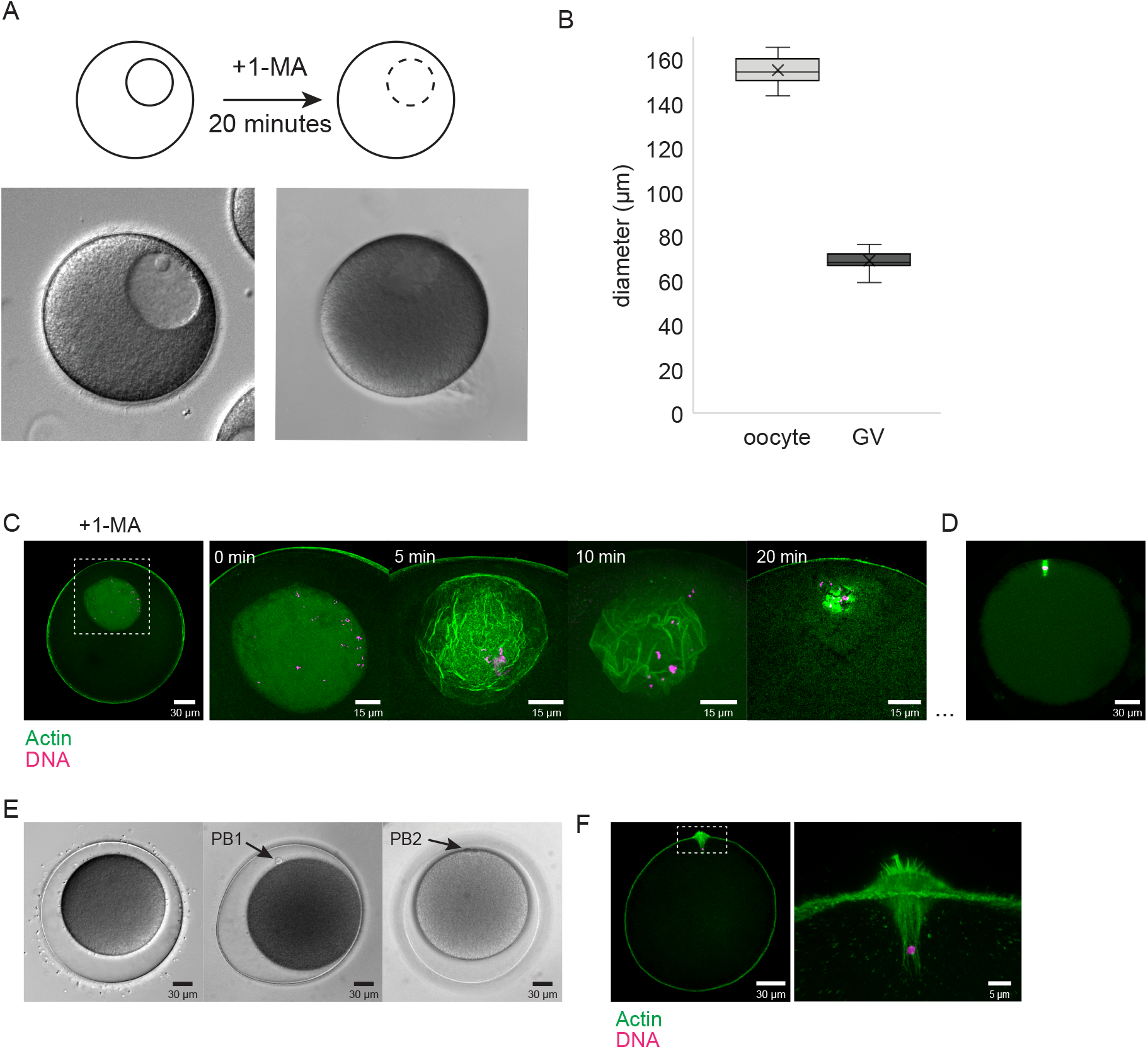
Cytoskeletal changes associated with *H. attenuata* oocyte maturation and fertilization. A) Schematic representation and corresponding DIC images of *H. attenuata* oocytes before and after 1-methyladenine (1-MA) treatment. B) Quantification of egg and germinal vesicle size before maturation. C) Representative confocal images of oocytes stained with phalloidin (actin, green) and Hoechst 33342 (DNA, magenta) before and after (5,10 and 20 minutes) treatment with 1-MA. D) Representative confocal image of 1-MA treated oocyte stained with Hoechst 33342 (DNA, magenta) and for tubulin (green). E) Representative brightfiled images of fertilized oocytes undergoing meiosis. PB= polar body. F) Representative confocal image of fertilized egg stained with phalloidin (actin, green) and Hoechst (DNA, magenta). All confocal images are projection of z-stacks covering the whole nucleus. All images are representative of a minimum of 3 repeats.

Like for other startfish species, meiotic resumpion could be induced *in vitro* by treatment with the hormone 1-methyladenine (1-MA, 10 µM) (Kanatani et al., 1969; Kishimoto and Kanatani 1976). Within 15 minutes of 1-MA addition (at 18 °C) the nuclear membrane become irregular and less defined and by 30 minutes of treatment, the germinal vesicle broke down (GVBD) and disappeared, indicating meiotic resumption (Figure 2A). Prior to GVBD, actin is enriched at the plasma membrane although a cytoplasmic actin filament network was also visible in GV stage oocytes (Figure 2C). This network disappeared in 1-MA treated eggs. At GVBD, a shell of actin polymerizes below the nuclear surface and contracts, bringing the chromosomes together (Figure 2C). The nucleolus which was present in the GV disappears at the end of nuclear breackdown. At this stage a small meiotic spindle, of about 10 µm in length, forms near the cell surface and chromosome can be seen aligned on the metaphase plate before anaphase (Figure 2D).

At 18 °C polar bodies were released approximately 50 and 80 minutes after 1-MA treatment (Figure 2E) both in the presence and in the absence of fertilization. Fertilization was performed by adding diluted sperm to the sea water containing the eggs. Within 1-2 minutes of sperm addition, the fertilization envelope rose from the surface of 95-100% of zygotes, indicating fertilization (Figure 2E). Eggs could be fertilized following GVBD and for at least 3 hours after 1-MA treatment. At fertilization actin filaments can be observed accumulating at the site of sperm entry (fertilization cone; Kyozuka and Osanai, 1988) and the male pronucleus can be observed associated with the microfilaments of the fertilization cone (Figure 2F). Following the release of the second polar body the female pronucleus, migrates from the animal pole towards the male pronucleus and pronuclear fusion takes place in the center, followed by mitosis and first cleavage.

### Embryonic development of *H. attenuata*

The cleavage divisions that follow fertilization are synchronous, holoblastic and equal (Figure 3). As in most animal embryos, the first cell division is slow and, at 18 °C, the first mitosis, measured as time from NEB (nuclear envelope breakdown) to cleavage, lasts 30 minutes while cytokinesis occurs 4 hours post fertilization (2-cell stage). The following divisions then occur every hour. The second and third divisions are radial. The second cleavage plane is meridian to the first while the third cleavage plane is perpendicular to the first two giving rise to a 2-tiered embryo with eight equal blastomeres (Figure 3D). The following divisions give rise to a spherical embryo with loosely connected blastomeres positioned underneath the fertilization envelope forming a hollow sphere (Figure 3F-I).

**Figure 3:**
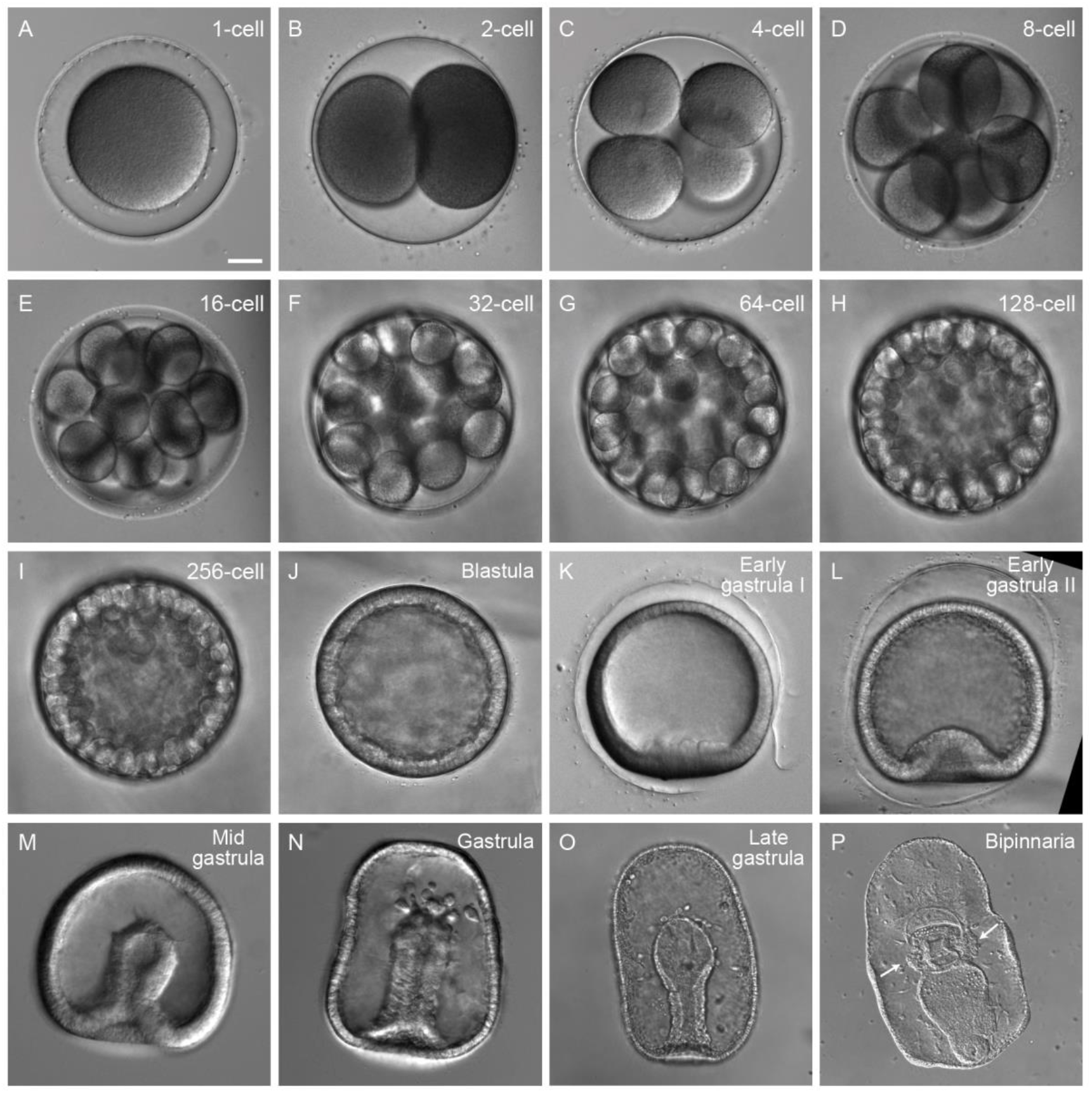
Timecourse of early embryonic development in *H. attenuata*. A-P) Brightfield images of *H. attenuata* embryos at different developmental stages from fertilization to early larva. Arrows in P point to coelomic pouches. Scale bar: 30 µm

About 20 hours post fertilization a blastula is formed. It consists of a single layer of packed cells enclosing a cavity, the blastocoel (Figure 3J). At this stage the embryo is covered in cilia and rotates within the fertilization envelope. After 24-26 hours the ventral side of the embryo thickens by elongation of cells in the vegetal side (Figure 3K). At this stage the fertilization envelope raptures and the embryo becomes a free-swimming blastula (Figure 3K). Mesenchimal cells can be observed budding off from the flattened ventral side and entering the blastocoel, indicating beginning of gastrulation (Figure 3K).

The embryo then undergoes embolic invagination and becomes a two layered gastrula (30 hpf). (Figure 3L). Gastrulation involves inward invagination on the ventral side resulting in the formation of a cavity, the archenteron, comunicating with the exterior by the blastopore (Figure 3M). During the invagination more mesenchymal cells can be observed delaminating from the tip of the growing archenteron and present within the embryo cavity (Figure 3N-O). At this stage the embryo elongates along the animal-vegetal axis and its entire surface is still covered by cilia (Figure 3P). Two coelomic pouches form on the anterior part of the invaginating archenteron at the bipinnaria stage (Figure 3P, arrows) which then will extend to give rise to the coelomic system.

### Larval development of *H. attenuata*

At 18 °C a bipinnaria larva develops in about a week (Figure 3P). It is a bilaterally symmetrical feeding larva of about 500 µm, with two ventral folds: a pre-oral and an anal hood (Figure 4). Six lateral lobes then develop, three on each side of the late bipinnaria larva. At this stage the archenteron has reached the anterior ectoderm and a complete, functional digestive system can be observed, with a mouth, a narrow proximal esophagus, a large round stomach (Figure 4 A-B’). From this stage onwords embryos were fed every day with a mix of unicellular algae (see material and methods).

**Figure 4:**
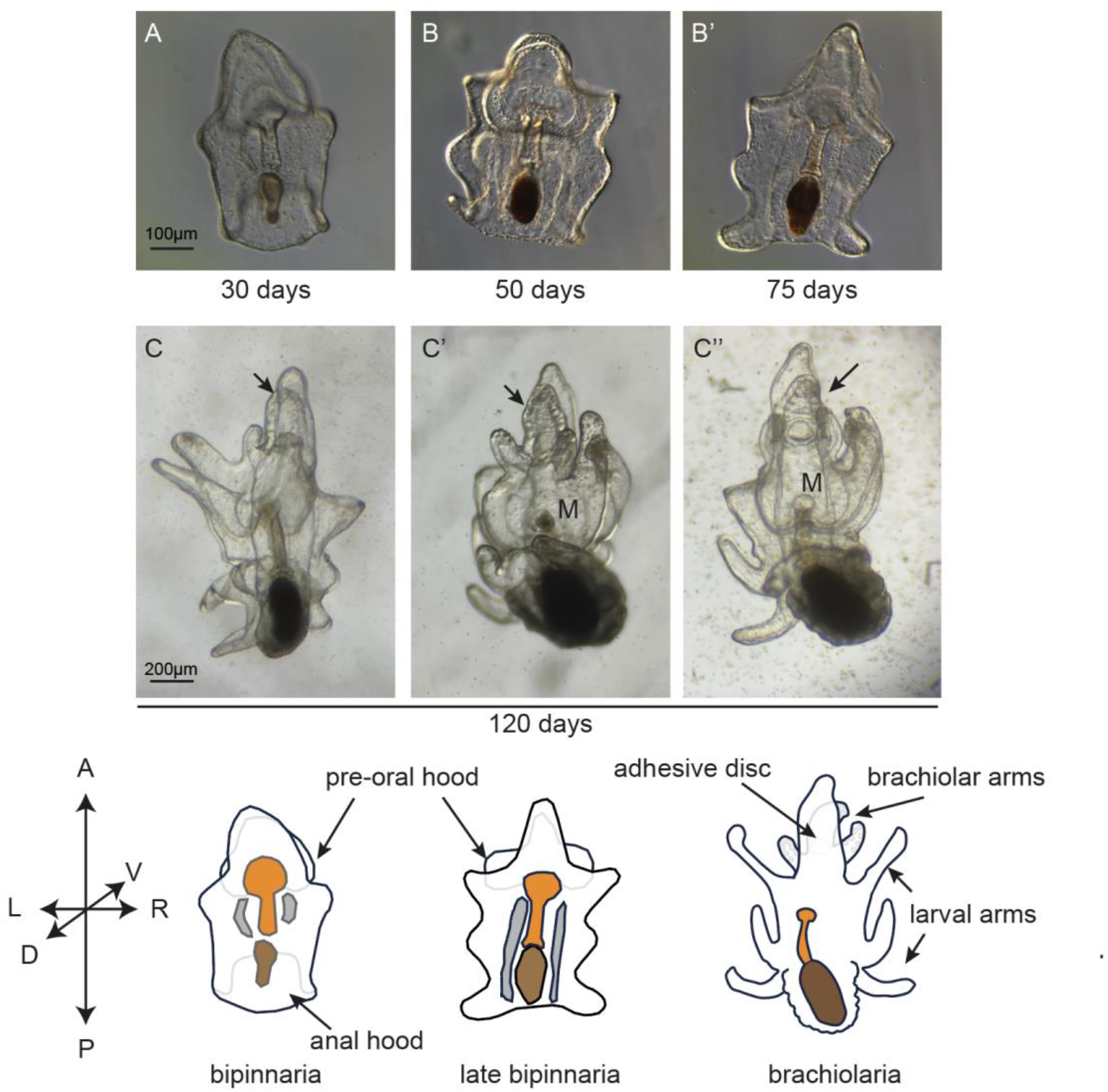
Larval development in *H. attenuata*. A-B’) Brightfield images of bipinnaria and C-C’’) brachiolaria larvae of H. attenuate. Arrows point to brachiolar arms. M= mouth D) Schematic representation of different larval stages with in esophagus in orange, stomach in brown and coelomic system in grey.

In the following 4-6 weeks the larvae grow to about 1-1.5 mm, giving rise to the brachiolaria larva. The side-lobes increase in length to become six long slender larval arms, which move and contract (Figure 4 C-C’’). At this stage, larvae respond to external stress by contracting their body and arms and relax in the presence of oxygen and food. At the brachiolaria stage, specialized attachment structures can be observed on the preoral lobe: the brachiolar arms, covered with papillae and the attachment disk (Figure 4C-C’’, 120 days, arrows).

### *H. attenuata* development at 23 °C

Recent work in another starfish species, *Pateria miniata* showed that an increase in water temperature of 4 °C significantly speeds up development at all stages (Barone et al., 2025). To determine the effect of temperature on *H. attenuata* development, we followed a similar protocol to that described for *P. miniata* and split siblings into three dishes and let them develop at 18 °C, 21 °C and 23 °C. The developmental times are reported in Table 1. As expected, development occurred faster as temperature increased. While at 18 °C cleavage divisions occurred every hour, at 23 °C blastomeres divided every 45 minutes and reached the blastula stage 14 hours post fertilization, 6 hours earlier than at 18 °C. The embryo then gastrulates and at 23 °C the archenteron reaches its full extension and expands at the tip by 48 hpf. Mesenchymal cells can be observed delaminating from the tip of the archenteron starting from 29 hpf and are present in the embryo cavity. The bipinnaria stage is reached after 60 hpf, 5 days faster than at 18 °C and then develops into brachiolaria larva within 20 days (20 days post fertilization, dpf).

**Table 1:** Developmental timining of *H. attenuata* at different temperatures. Time 0 is extrusion of the second polar body. For reference, the timing of equivalent stages of *P. miniata* embryos is also shown (taken from Barone et al., 2025).

|  | *Hacelia attenuata* |  |  | *Patiria miniata* |  |
| --- | --- | --- | --- | --- | --- |
|  | 18°C | 21°C | 23°C | 16°C | 20°C |
| 2cell | 2h40' | 2h10' | 1h40' | 2h30' | 2h |
| 4cell | 3h50' | 3h | 2h25' | 3h15' | 2h30' |
| 8cell | 4h50' | 3h50' | 3h10' | 4h15' | 3h10' |
| 16cell | 5h50' | 4h40' | 4hr | 5h | 3h50' |
| 32 cell | 6h50' | 5h30' | 4h45' | 5h45' | 4h20' |
| 64-cell | 7h50' | 6h20' | - | 6h45' | 5h |
| 128-cell | 8h50' | 7h10' | - | 7h30' | 5h40' |
| 256-cell | 10h20' | 8h10' | - | 8h15' | 6h15' |
| Blastula | 20h | 15h | 14h | 18h | 12h |
| Ventral thickening | 24-26h | 20h | 18h |  |  |
| Early gastrula | 30h | 22h | 22h | 24h | 16h |
| Mid-gastrula | 48h |  | 26h | 27h | 18h |
| gastrula | 72h |  | 29h | 34h | 24h |
| Late gastrula |  |  | 48h | 48h | 30h |
| bipinnaria | 7d |  | 60h | 72h | 48h |

## Discussion

This study provides the first complete description of the development of the Mediterranean starfish *Hacelia attenuate*, from oocyte maturation and fertilization through embryogenesis to larval formation.

Most starfish species have a single reproductive season and are dioecious free-spawners with a biphasic life cycle having a free-swimming planktonic larva which metamorphoses into a benthic, reproductively active adult (Jagersten, 1972). A minority of species instead deposit eggs on the substratum, where they undergo benthic development (Chia, 1968; Byrne, 1995), or retain their eggs in brood chambers (McEdward and Miner, 2001). Our data place H. attenuate within the first group.

Five larval forms have been described in the class Asteroidea—bipinnaria, brachiolaria, yolky brachiolaria, barrel-shaped larvae and yolky non-brachiolaria larvae (Fell, 1967; Oguro, 1989; Chia et al., 1993; McEdward and Janies, 1993)—and our data place *H. attenuata* within the most widespread canonical bipinnaria–brachiolaria types of larvae. The bipinnaria of *H. attenuata* is a bilaterally symmetrical, ciliated larva with hollow arms lacking skeletal rods, consistent with descriptions from other species (MacBride, 1914; Kume and Dan, 1968). At the brachiolaria stage we observed the characteristic attachment structures on the preoral lobe—brachiolar arms covered in papillae and a distinct attachment disc— usually used to probe the substratum, sense metamorphic cues and adhere during settlement (Haesaerts et al., 2006; Murabe et al., 2007). Thus, although *H. attenuata* occupies a distinct biogeographic location, its overall life cycle and larval morphology are highly conserved relative to other free-spawning starfish.

Our observations of cytoskeletal dynamics during meiosis and fertilization reveal striking parallels with other starfish species and highlight conserved mechanisms of chromosome capture. Meiotic resumption in *H. attenuata* can be reliably induced *in vitro* by 1-methyladenine, as for other laboratory starfish species. We observed a cytoplasmic actin network expanding the whole oocyte in prophase-I oocytes that disappeared upon hormone treatment and the formation of an actin shell within the germinal vesicle at GVBD. Contraction of this shell then gathered dispersed chromosomes toward the future spindle region, similar to the actin-dependent chromosome-capture mechanism described for *Patiria miniata* oocytes (Lenart et al., 2005). Fertilization in *H. attenuata* was accompanied by the rapid formation of a fertilization envelope and a fertilization cone at the sperm entry site, with the male pronucleus remaining associated with actin filaments before it migrated toward the cell center. The morphology and timing of this structure closely resemble the fertilization cone previously characterized in the Mediterranean starfish *Astropecten aranciacus* (Chun et al., 2010). Together, these cytological observations underscore the value of *H. attenuata* as a comparative model for dissecting the interplay between actin dynamics, spindle assembly and pronuclear migration in large marine oocytes. For exemple it will be interesting to determine if sperm centration occurs during interphase like the sea urchin *Paracentrotus lividus* (echinorderm, Tanimoto et al., 2016) or rather at mitotic entry like the ascidian *Phallusia mammillata* (tunicate, Rosfelter et al., 2024)

Early embryogenesis in *H. attenuata* also conforms broadly to the stereotyped radial cleavage pattern of echinoderms. Cleavages were equal and holoblastic, with meridional first and second divisions and an equatorial third division producing an eight-cell, two-tiered embryo. Subsequent divisions yielded a loosely cohesive embryo under the fertilization envelope; removal of this envelope prevented the formation of cell–cell contacts, highlighting its mechanical role in maintaining early embryo integrity. Gastrulation proceeded by invagination at the vegetal (future ventral) pole, with delamination of mesenchymal cells from the archenteron tip and the early appearance of paired coelomic pouches (Perillo et al., 2023). These features confirm that *H. attenuata* follows the canonical starfish mode of embryo development while providing precise staging criteria for future molecular and functional studies.

The geographic and ecological context of *H. attenuata* is particularly relevant for interpreting its developmental traits. Originally described by Gray in 1840, *H. attenuata* is distributed throughout the Mediterranean Sea at depths ranging from 1 to 150 m, but is most frequently encountered below 20 m (Hansson, 1999). Additional records from the Azores (Micael et al., 2012), Canary Islands, Cape Verde and the Gulf of Guinea (Hansson, 1999) indicate that it spans warm-temperate to subtropical environments. Morphologically, the species belongs to the order Valvatida and is easily recognized by the longitudinal rows of papulae along the arms. Our data on temperature-dependent development fit well with this biogeographic pattern. We found that raising rearing temperature from 18°C to 23°C accelerated all stages of development, from cleavage and blastula formation to gastrulation and the bipinnaria and brachiolaria stages, without obvious defects in morphology. Embryos at 23°C reached key milestones several hours to days earlier than siblings at 18°C, similar to the pattern reported for *P. miniata* at temperatures 4°C higher than the optimal for that species (Barone et al., 2025). Preliminary observations of *H. attenuata* larval survival and metamorphosis suggest that developmental success remains high across this temperature range. Taken together, these findings indicate that *H. attenuata* is well adapted to relatively high temperatures, consistent with its distribution in the warm, saline waters of the Mediterranean and adjacent Atlantic regions.

From a broader ecological and evolutionary perspective, our work expands the limited set of echinoderm and, in particular, starfish species for which complete developmental series are available. Because *H. attenuata* combines a typical indirect life cycle with robust development at elevated temperatures, it offers a promising resource for future studies on thermal tolerance, developmental plasticity and the molecular basis of adaptation to high-salinity conditions. Comparative analyses with established model species such as *Patiria* and *Asterias* will help disentangle conserved from lineage-specific developmental programs and may reveal how subtle changes in early development influence the resilience of Mediterranean benthic communities. This combination of phylogenetic comparability and environmental relevance also establishes *H. attenuata* as a powerful new model for investigating how climate-driven changes in ocean conditions may impact starfish development, population dynamics and ecosystem roles.

## Acknowledgments

We are grateful to L. Besnardeau for technical support and to A. McDougall and R. Dumollard for sharing of resources. We thank the imaging platform, PIM (member of the MICA microscopy platform) where image acquisition was conducted, and the CRB (Center for Biological Resources) for support with animal collection and housing. Both PIM and CRB are supported by EMBRC-France, whose French state funds are managed by the ANR within the investments for the future program under reference ANR-10-INSB-02.

